# Estrogen and progesterone exhibit distinct yet coordinated roles in the regulation of tendon extracellular matrix remodeling

**DOI:** 10.1101/2025.03.05.641754

**Authors:** Allison M. Sander, Brianne K. Connizzo

## Abstract

Remodeling of the extracellular matrix (ECM) is required for the proper healing, strengthening, and maintenance of tendon tissue. There are well documented sex differences in tendon injury rates and healing outcomes, often attributed to either innate differences in tissue structure and resident cell signaling or the influence of sex hormones. However, these factors are rarely decoupled. Estrogen (17β-estradiol) and progesterone (P4) receptors are expressed in both male and female tendons and thus could participate in the remodeling process, but studies are extremely limited. Therefore, the objective of this work was to address whether biological sex differences are present in tendon remodeling and to determine the individual and combined roles of estrogen and progesterone in the remodeling process. We utilized an explant model of the flexor digitorum longus tendon harvested from young adult male and female mice to examine cell-mediated remodeling without disruption to the native environment. We found sex differences in tendon remodeling in the absence of hormonal stimulation, revealing how biological sex alone influences tendon health. We also demonstrate that the response to exogenous hormone delivery is sex-dependent, and that progesterone and estrogen serve complimentary yet independent roles. Overall, this work presents the first examination of sex-dependent matrix turnover in response to hormones and underscores the critical need for additional research in this area.

## Introduction

Sexual dimorphism in tendon injuries and healing outcomes are well recognized, yet not well understood. One study reports an almost 10% increase in rotator cuff pathology per person years in females compared to males,^1^ and it is well known that anterior cruciate ligament (ACL) injuries have 3 to 6 times higher prevalence in young females directly following the onset of puberty.^2^ Interestingly, Achilles tendon rupture is 5 to 6 times more prevalent in active middle-aged men compared to age-matched women.^3^ Despite the NIH mandate on considering sex as a biological variable, much of the research still only focuses on male samples.^4–6^ In fact, a systematic review found that only 2% of the total participants represented in the literature on patellar tendinopathy are female, and just 5% recognized the lack of female representation as a limitation of their work.^4^ Due to the deficit of female data, it is difficult to understand the sex differences that exist or the underlying factors which contribute to them.

To investigate injury prevalence, sex differences in tissue mechanical function have been investigated in both human studies and in pre-clinical animal models.^7^ For example, it was shown that inactive males had significantly longer and thicker Achilles tendons than age- and activity-matched females, resulting in increased *in vivo* peak stresses.^8^ Data in murine tendons conflict with these findings, with decreased mechanical properties in male Achilles and supraspinatus^9,10^ tendons compared to females. Studies on cadaveric Achilles tendons could also not replicate this finding, demonstrating no significant differences in mechanical properties between sexes.^11^ The discrepancies between studies, especially comparing humans to rodents, suggest that sex-related changes may be a result of epigenetic history or specific site of the tendon. Therefore, active cell-mediated processes, such as injury and recovery, may hold more answers than structure-function studies alone. Injury and healing studies have indeed demonstrated significant sex-related differences. After ACL transection, female minipigs had significantly diminished healing outcomes in all measured parameters of graft structural properties, knee laxity, and cartilage damage compared to males.^11^ These structural sex differences were also replicated in a murine ACL injury model.^12^ While this supports the notion that there are programmatic sex differences in tissue remodeling, healing involves significant immune responses which could cloud our view of tissue-specific phenomena.

We recently established a murine explant model of extracellular matrix remodeling in the absence of a macroscale injury, induced by changes in mechanical stimulus through either stress deprivation^13^ or increased mechanical loading.^14^ This model is advantageous as the effects of innate genetic programming differences and the influence of circulating or local factors, such as sex hormones or inflammation, can be decoupled. Our previous work revealed innate sex differences in metabolic activity and cell proliferation, and hinted at altered synthesis of extracellular matrix components.^13^ However, remodeling is a delicate, multi-step and multi-scale process, involving clearance of damaged ECM, synthesis of new ECM, incorporation and repair of existing ECM proteins, and reorganization of matrix structure. Therefore, a more complete characterization is necessary to fully understand sex-specific remodeling processes. Furthermore, our previous work did not consider the presence or absence of sex hormone signaling, a factor likely to be important to female tendon health and homeostasis.

Changes in hormone signaling, such as post-menopausal drops in the gonadotropic female-associated steroid hormones estrogen and progesterone, are associated with many health concerns, including musculoskeletal pathologies.^15–17^ Estrogen and progesterone can interact with DNA directly or with targets in the cytosol to alter transcription and ultimately protein synthesis.^18–21^ The presence of estrogen (17β-estradiol) and progesterone (P4) receptors in resident tendon cells,^22^ tenocytes, suggest sex hormones could directly influence tendon biology yet their role in ECM turnover is still not well understood. A large majority of the studies investigating hormone effects on tendon health have focused on the influence of estrogen on ECM synthesis. Using serum levels of hormone for comparison between groups, clinical studies have connected increased estradiol levels with decreased response to mechanical loading,^7,15,23–25^ and higher tendon fractional synthesis rates.^15^ However, there are a number of confounding variables in human studies, such as the type of estrogens studied, the level of activity of participants, and the lack of reported chemical makeups of oral contraceptives used by participants, which could all influence the impact of hormones on tendon remodeling.^23,25,26^ In rodent studies, estrogen has been shown to increase fibroblast proliferation and promote relaxin receptor expression,^27–30^ which can lead to increased breakdown of collagen. Supporting this, estrogen-deficient states have been linked to decreases in tenocyte proliferation and metabolism^27,31^ and inferior healing outcomes.^32–34^ This indicates estrogen may have complex interactions with tenocytes which influence both the breakdown and synthesis of tendon ECM.

Progesterone, on the other hand, has not been deeply studied in tendon. The effects of progesterone have also not been decoupled from estrogen as the human menstrual cycle does not have a phase in which progesterone levels are isolated from estrogen. However, both hormones together (post-ovulatory phase of the menstrual cycle) have been shown to increase collagen synthesis more than estrogen alone (pre-ovulatory phase).^35^ Progesterone receptors have sex-dependent binding mechanisms, implying progesterone could be a key component in the explanation of sex differences.^36^ However, the musculoskeletal literature hasn’t been clear on whether progesterone induces anabolic or catabolic signaling. In rodents, progesterone has also been linked to increased relaxin receptor expression, stimulating collagen breakdown.^30^ Conversely, progesterone has been linked with increased collagen synthesis rates in muscle from post-menopausal women treated with the exogenous hormones^37^ and administration of progesterone to male mice with induced myocardial infarctions *in vivo* revealed progesterone can promote cardiac tissue maturation.^38^ Progesterone is known to play a key role in the maturation of fetal tissues, through development of both the fetal immune^39^ and central nervous systems.^40^ In addition, pelvic floor prolapse has been highly correlated with dysfunctional progesterone receptors.^41^ Given the similarities in collagen structure between pelvic floor muscle and tendon, these data suggest an important role for progesterone in extracellular matrix maturation and turnover that has not yet been examined.

The overarching objective of our studies was to determine the roles of sex and sex hormones in ECM remodeling following stress deprivation. Using our explant model, we aimed to disentangle the independent effects of biological sex, estrogen, and progesterone on the synthesis and breakdown of key extracellular matrix molecules. Based on the published literature and preliminary data from the lab, we hypothesized that sex hormones are a main contributor to sex differences in tendon remodeling, with estrogen being associated with ECM turnover and progesterone contributing to ECM maturation. We also expected the influence of the combined hormones to be additive based on previous clinical data.

## Materials and Methods

### Sample Preparation

We sought to explore responses to hormones in both male and female tissues in this study as many previous rodent studies were only performed in males, ^28,33,42,43^ assuming the response would be comparable in female tissues. Male and female young adult (4 month; n = 120 animals) C57B/6J mice were housed in the animal facility for at least one week to mitigate impacts from hormonal fluctuations due to the stress of travel.^44–47^ Flexor digitorum longus (FDL) tendon explants were then harvested in a biosafety cabinet (BSC) as previously described^48^ per approved animal use protocol (BU IACUC PROTO202200000074). Harvested tendons were washed twice in sterile PBS and 100μg/mL streptomycin (Fisher Scientific), and 0.25μg/mL Amphotericin B (Sigma-Aldrich) before being introduced to culture conditions.

### Culture Conditions

Explants were cultured in stress-deprived conditions (no mechanical stimulation) for up to seven days or collected immediately to assess baseline properties (Figure 1). The incubators maintained a constant environment of 37°C and 5% CO_2_. The culture medium base consisted of Dulbecco’s Modified Eagle’s Media (1g/L; Fisher Scientific, ‘DMEM’) supplemented with 10% fetal bovine serum (Cytiva, ‘FBS’), 100units/mL penicillin G, 100μg/mL streptomycin (Fisher Scientific), and 0.25μg/mL Amphotericin B (Sigma-Aldrich). There were four medium groups: no hormone (base medium only, ‘NH’), estrogen-supplemented (base + 1nM water-soluble 17β-estradiol suspended in 0.1% BSA, E4389, Sigma-Aldrich, ‘EST’),^42,49–52^ progesterone-supplemented (base + 1nM progesterone suspended in 0.1% BSA, P7556, Sigma-Aldrich, ‘PRO’),^50–53^ and dual-hormone-supplemented (base + 1nM water-soluble 17β-estradiol + 1nM progesterone, ‘DH’). All hormones exogenously added were chemically equivalent to the endogenous hormones present in mouse serum. Medium changes occurred every other day and spent media were collected on day 6 (D6) to assess matrix metalloproteinase (MMP) activity. Explants were collected at D0, D3, and D7 to assess metabolic activity, matrix biosynthesis, composition, and gene expression.

**Figure 1.**
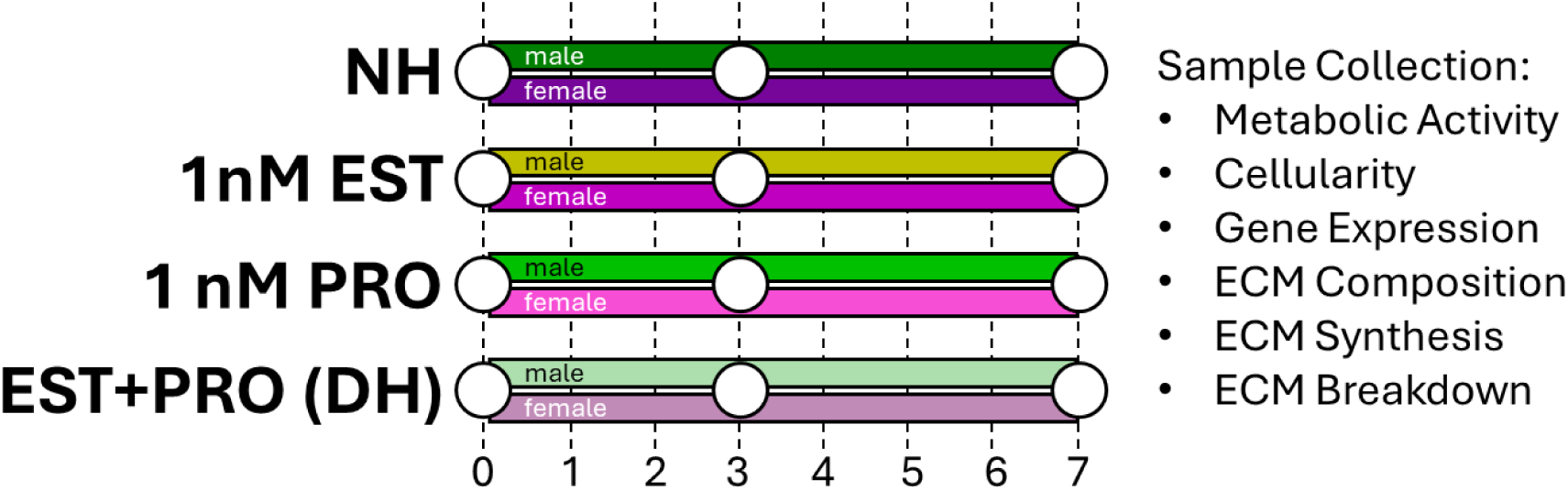
Overall study design schematic depicting eight groups that were studied: no hormone (‘NH’), estradiol-treated (‘EST’), progesterone-treated (‘PRO’), and dual hormone treated (‘DH’) for both sexes. Explants were assessed for changes in metabolic activity, cellularity, gene expression and ECM composition at baseline (day 0) and after 3 and 7 days in culture. Measures of real-time ECM synthesis and breakdown were also quantified at days 3 and 7.

### Matrix Biosynthesis, Explant Composition, Metabolism, and DNA Content

Radioactive isotope labels ^35^S-sulfate (2μCi/mL) and ^3^H-proline (1μCi/mL) were added for 24 hours on the last day of culture to measure the synthesis of sulfated glycosaminoglycans (GAG) and total protein, respectively (Revvity). One well of media without an explant was used as an internal control. For the last 3-hours of culture, resazurin solution (10% v/v) was added to the media to assess explant metabolism as previously described.^54^ Fluorescence intensity of the product, resorufin, was measured in collected medium following culture at 554/584nm, and sample wells were normalized to the values from the control (medium-alone) well. Further, D3 and D7 metabolism were normalized to D0 readings from samples of the same sex and hormone treatment, such that a value of 1 would indicate D0 levels of metabolic activity. This normalization was performed as differences were seen after just 3 hours of hormone stimulation between study groups. Explants were then stored at −20°C until further processing.

Wet weights were taken by hydrating explants in PBS for 1 minute, dabbing on a paper towel, and weighing. Explants were lyophilized for at least 3 hours, and dry weights for each sample were then measured. All weights were performed in triplicate and averaged. Explants were then digested using 5mg/mL Proteinase K (Sigma Aldrich). Sample digests were assessed for protein synthesis and GAG synthesis via radiolabel incorporation rate using a scintillation counter (Perkin Elmer). DNA content was determined using a PicoGreen assay (Quant-IT DNA).^55^ Total collagen and GAG content were assessed through hydroxyproline (OHP) and dimethylmethylene blue (DMMB) assays respectively.^56,57^ All measures were normalized to sample dry weight.

### Matrix Degradation

Spent culture media collected at day 6 was stored at −20°C until use. Pooled activity of MMPs 1, 2, 3, 7, 8, 9, 10, 13, and 14, indicative of matrix degradation in tendon, was determined through analysis of spent culture medium (n=5/media type/sex) using a FRET-based generic MMP activity kit (SensoLyte® 520 Generic MMP Activity Kit Fluorimetric; AnaSpec). This assay measures generic activity by introducing an MMP substrate that fluoresces when cleaved by MMPs in the medium sample. Relative cleavage activity is measured via fluorescence intensity at 490/520nm.

### Gene Expression

Baseline (D0) stress-deprived samples were washed in PBS and 100units/mL penicillin G, 100μg/mL streptomycin (Fisher Scientific), and 0.25μg/mL Amphotericin B (Sigma-Aldrich) solution and immediately flash frozen in liquid nitrogen. Cultured samples were washed in PBS for 1 minute before flash freezing to remove any culture media. Samples were stored at −80°C until use. On the day of RNA isolation, explants were submerged in Trizol and homogenized with a bead homogenizer (Benchmark Scientific). The sample supernatant was separated using phase-gel tubes (Qiagen) and purified using a Zymo Quick-RNA purification kit (Zymo). RNA concentration of each sample was quantified using an Agilent BioTek Take3 Microvolume Plate (Agilent). These concentrations were used to calculate dilution volumes such that each sample had the same amount of RNA loaded for reverse transcription (RT). After RT, qPCR was conducted on cDNA samples using Applied Biosystems StepOne Plus RT-PCR (Applied Biosystems). Results were analyzed using the ΔΔCT method and were normalized to the housekeeping gene TBP (TATA box binding protein)^58–60^ and to D0 samples of the same sex, such that a value of 0 would indicate innate expression of the gene.

### Statistical Evaluation

Statistical evaluation was performed on the no hormone groups for innate sex comparisons through two-way ANOVAs with Tukey’s multiple comparisons tests where appropriate based on passage of Brown-Forsythe tests. Within day and sex groups, one-way ANOVAs with Tukey’s multiple comparisons tests where appropriate were used to evaluate the response to hormonal treatments. In the comparisons between hormonal treatments and all qPCR analyses, two-tailed t-tests were used to determine differences from day 0 measurements. Significance is reported at p<0.05.

## Results

We first assessed the impact of biological sex and hormones on explant health. In the absence of hormones, sex differences were present in metabolic activity and dry weight, with females having lower values compared to males (Fig. 2B,D), aligning with the increased size of male tendon tissues measured previously.^7–9^ After just 3 hours of hormone induction, DNA content for both male and female groups dropped, as shown by the decreased baseline levels (Fig. 2A). Progesterone caused a further decrease in DNA content from D0 in males. Interestingly, estrogen alone maintained female NH levels of DNA content, but when in combination with progesterone, content was decreased from both D7 NH and D0 levels. Male sensitivity to progesterone is also seen in metabolism, as this is the only treatment which increased metabolic rates above both D0 and NH male levels (Fig. 2B). In females, all hormone treatments led to increased metabolic activity levels which mimicked that of males. Both hydration and explant dry weights remained relatively constant throughout culture (Fig. 2C-D). The dual hormone condition, however, led to increases in hydration from D0 in both groups.

**Figure 2.**
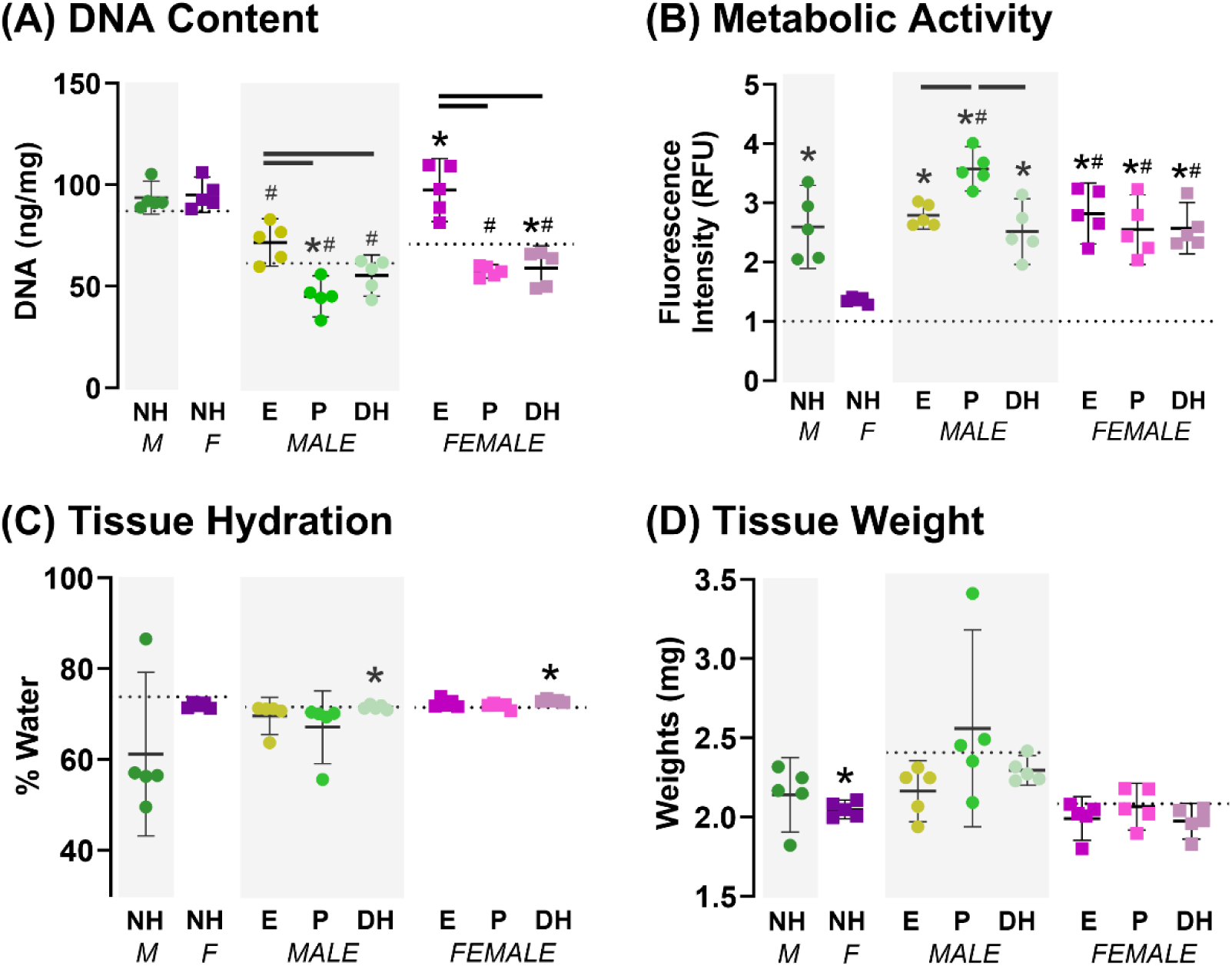
Impacts of sex and sex hormones on explant health at D7. For each graph, male (M) and female (F) no hormone (NH) groups are compared on the left, the effects of hormone treatment (EST, PRO and DH) on males in the center panel, and the impact of hormonal treatment on females on the right. Health parameters shown are DNA content (A ), relative metabolic activity (B) which is normalized to D0 activity, explant hydration (C), and dry weights (D). All data are represented as mean ± 95% confidence interval, with solid lines representing significant differences between groups, dashed and dotted lines representing D0 values, * representing differences from D0, and # representing differences from the NH treatment of the same sex, all reported at p<0.05. Male data is highlighted with grey background while female is presented with white background.

We then isolated the influence of biological sex alone on tendon remodeling through comparisons groups with no hormone treatment (Fig. 3). In both males and females, overall collagen content and collagen 1 expression increased throughout the culture period (Fig. 3A-B). However, while the males exhibited increases in protein synthesis rates from D3 to D7, female protein synthesis remained constant (Fig. 3C). MMP activity remained constant in both males and females at similar rates (Fig. 3D). However, when examining MMP-9 and MMP-13, which specifically degrade collagen, both males and females show increases in expression from D3 to D7, with males having greater expression than females at D7 (Fig. 3E-F). When looking at non-collagenous matrix turnover, males exhibited an overaccumulation of GAGs during culture while GAG content was maintained in female tissues (Fig. 3G). However, GAG synthesis rates were equivalent between the sexes at both measured timepoints and increased from D3 to D7 (Fig. 3H). Females generally had lower proteoglycan expression compared to males (Fig. 3I-K), especially by D7, where males had increased biglycan expression from innate levels. Expression of MMP-3, which specifically degrades GAGs and proteoglycans, was higher than baseline levels in both sexes but males had greater relative expression (Fig. 3L).

**Figure 3.**
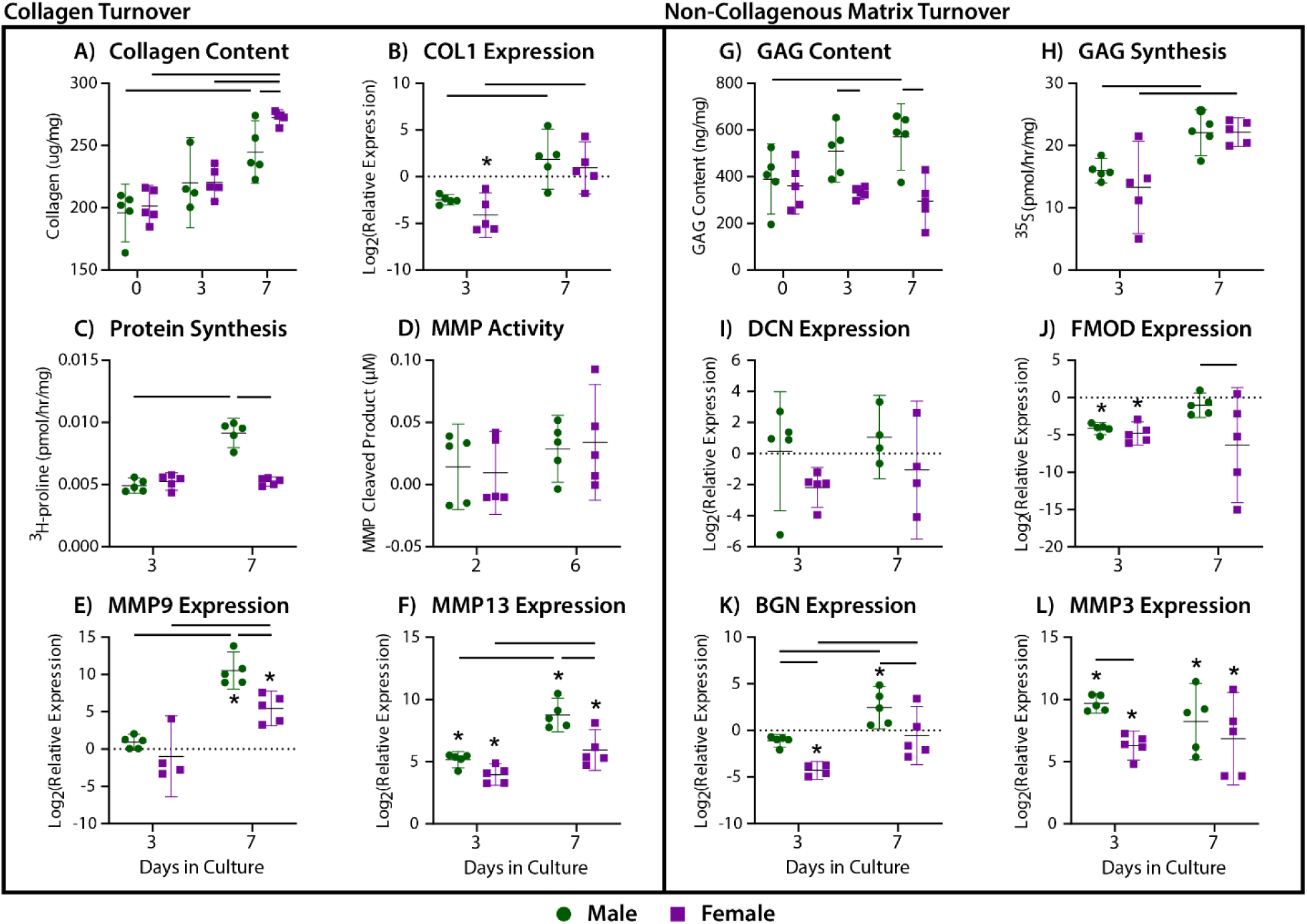
(Left) Measures of collagen synthesis, such as (A) collagen content, (B) gene expression of collagen 1, and (C) proline incorporation, and measures of collagen breakdown, such as (D) enzymatic MMP activity, and gene expression of (E) MMP-9 and (F) MMP-13, comparing males (green) and females (purple) in the no hormone conditions. (Right) Non-collagenous matrix turnover, measured by (G) sGAG content, (H) sGAG synthesis, and gene expression of (I) decorin, (J) fibromodulin, (K) biglycan, and (L) MMP-3. Data is presented as individual values with mean and 95% confidence interval noted. Statistical comparisons (p<0.05) between timepoints are represented by bars spanning between data points, while stars (*) represent differences from baseline d0 values

We then investigated the impact of the individual and combined hormones on collagenous matrix turnover. By D7, the single hormone conditions returned collagen content back to innate (D0) levels in both sexes while the dual hormone induction mimicked the no hormone group, with accumulation of collagen in both timepoints (Fig. 4A, G). Collagen 1 expression was maintained in males with the single hormone additions, but the dual hormone addition led to a significant decrease at D7 (Fig. 4B). In females, hormones led to trending decreases at both timepoints in collagen 1 expression, with a significant decrease with the dual hormone condition as well at D7 (Fig. 4H). Looking at proline incorporation, indicative of collagen synthesis, we see that the progesterone and dual hormone additions led to decreased rates at D3 in both sexes (Fig. 4C,I). By D7, single hormone groups in males have synthesis rates similar to the no hormone condition, but the dual condition exhibited a significant decrease. In females, the single hormones led to significant increases in synthesis rates, surpassing that of NH males even, while the dual hormone condition maintained the NH female level. Examining tissue breakdown, we saw that hormones had an additive impact, increasing MMP activity throughout culture compared to the baselines of both sexes, especially with the dual hormone condition (Fig. 4D, J). By D7, MMP-9 and MMP-13 specifically were maintained with no significant differences from the NH group of each sex with any hormone conditions and were all elevated from D0 levels (Fig. 4E-F, K, L).

**Figure 4.**
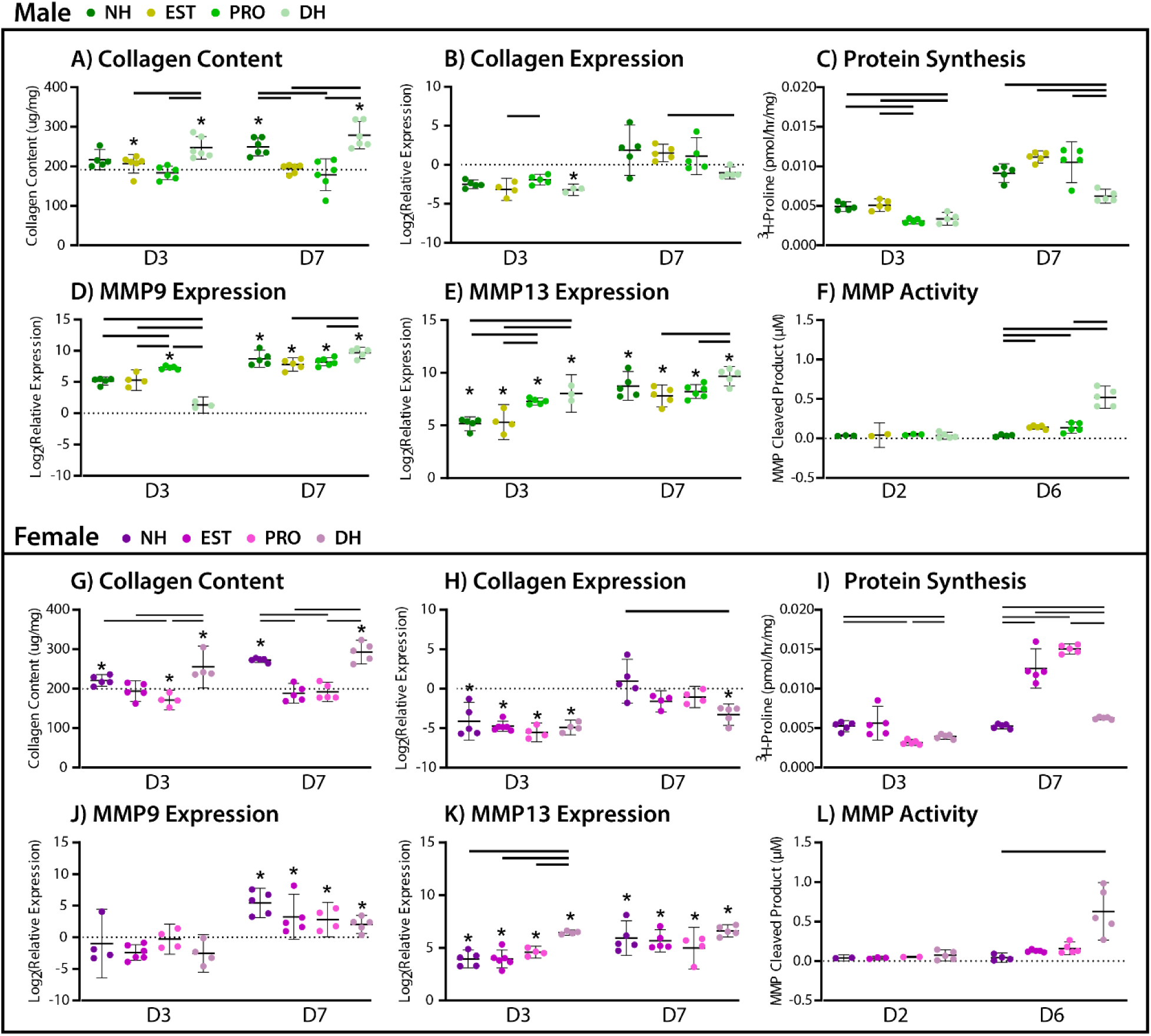
Impact of hormones on collagen turnover in male (top) and female explants (bottom) at day 3 and day 7 with no hormone (NH), estrogen (EST), progesterone (PRO) and dual hormone groups for each day. Collagenous matrix content (A,G), collagen 1 expression (B,H), overall protein synthesis (C, I) MMP9 (D,J) and MMP13 (E,K) expression, and overall MMP activity at day 2 and day 6 (F,L). Statistical comparisons (p<0.05) between hormone groups are represented by bars spanning between data points, while stars (*) represent differences from baseline d0 values.

Finally, we examined the influence of hormones on non-collagenous matrix turnover (Figure 4). While hormone addition did not lead to any differences in males, the progesterone and DH groups were best at maintaining innate (D0) GAG content (Fig. 5A). In females, estrogen led to a unique overaccumulation of GAGs by D7 while progesterone led to a decrease, adding together in the dual hormone condition to maintain baseline GAG content (Fig. 5G). Across sexes, the dual hormone condition generally led to a decrease in GAG synthesis from the no hormone baseline, again being an additive effect of the single hormone conditions (Fig. 5B, H). Proteoglycan expression was largely maintained at NH levels across sexes with the addition of hormones (Fig. 5C-E, I-K). However, at D3, progesterone supplementation led to decreases in female proteoglycan expression (decorin, fibromodulin) and at D7, the dual hormones led to proteoglycan expression which dipped below innate (D0) levels. While still maintaining increased expression from innate levels, progesterone led to a decrease in MMP-3 expression for males and an increase for females at D3 from their respective NH groups (Fig. 5F, L).

**Figure 5.**
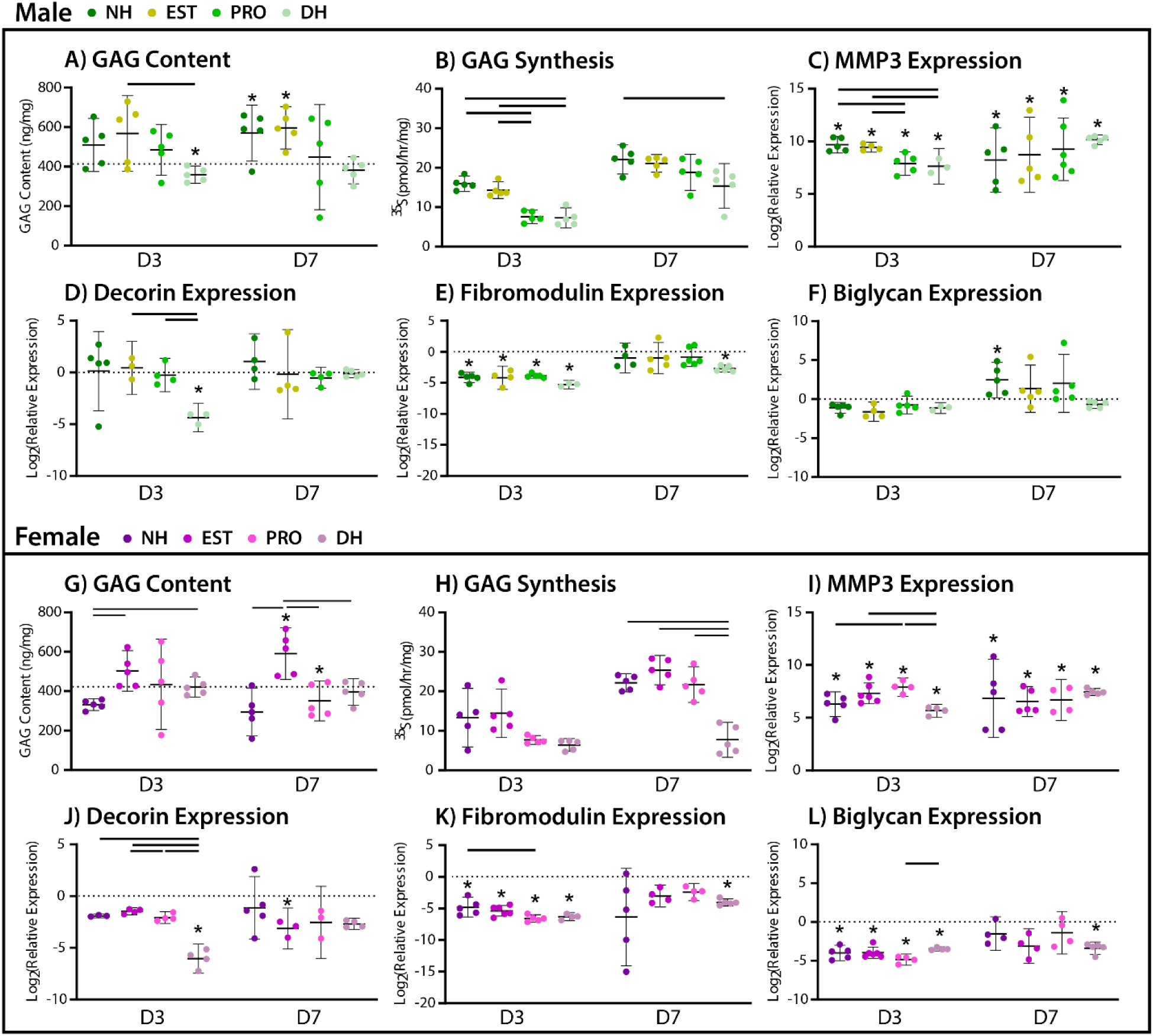
Impact of hormones on non-collagenous matrix turnover in male (top) and female explants (bottom) at day 3 and day 7 with no hormone (NH), estrogen (EST), progesterone (PRO) and dual hormone groups for each day. GAG content (A,G), synthesis (B,H), and MMP3 (C, I) decorin (D,J) fibromodulin (E,K), and biglycan expression (F,L). Statistical comparisons (p<0.05) between hormone groups are represented by bars spanning between data points, while stars (*) represent differences from baseline d0 values.

## Discussion

Consistent with our previous work, we found innate sex differences in many measured biosynthetic parameters. This could indicate possible chromosomal differences between male and female tendon programming that could impact their responses to a variety of changing stimuli (injury, aging, disease, etc.). In the context of explant health, we found increased dry weight in male tendons, which is consistent with previous work reporting increased length and CSA of male tendons.^13^ Increased metabolic rate in males was also replicated from our previous work^13^ and supports the conclusion that, when under the injury stimulus of stress-deprivation, male tendons rebuild while female tendons preserve matrix in the absence of hormonal stimulation. This is also supported by the findings of decreased protein synthesis, proteoglycan gene expression, and MMP gene expression in females compared to males. Connecting back to previous remodeling literature, this lack of remodeling response may also explain the decreased remodeling in female tendons in response to exercise^7^ and decreased healing outcomes^5^ when compared to males at the same timeframe. Our data therefore show that differences rooted in biological sex contribute to the decreased remodeling response of females, possibly acting to preserve matrix in response to injurious stimulus. Future studies will explore biological sex differences through more specific genetic approaches, isolating the influence of the hormonal environment from those of chromosomal factors with chromosomal invariant mouse models.^61^

When adding estrogen to female tissues, we see more similarities to the male tissue than to the female NH condition (Fig. 6). This indicates that estrogen supplementation may compensate for or reverse some of the innate sex differences we saw in tendon and improve the response of female tendons to injury. Furthermore, the bulk increase in female metabolism in response to estrogen seems to indicate that hormones are important for healthy maintenance of cell activity. Our data imply estrogen has a role in the stimulation of matrix turnover, increasing synthesis and degradation parameters in female tendon explants. This aligns with previous findings that estrogen bolsters cell proliferation in female musculoskeletal tissue.^27,28,34^ Turnover of non-collagenous proteins is known to be much faster than turnover of the collagen matrix,^62^ and could have a stronger impact on tendon function. Interestingly, we did find decreased DCN and FMOD expression from innate levels in females when estrogen is present. These proteoglycans are well known for regulating fibril structure during fibrillogenesis and healing, so it’s not clear whether their decreased expression suggests altered non-collagenous turnover or altered collagenous organization. Thus, it will be important to include more timepoints, different types of injury, and a more specific proteomic assessment of GAGs and proteoglycans to make an accurate assessment of estrogen’s role.

**Figure 6.**
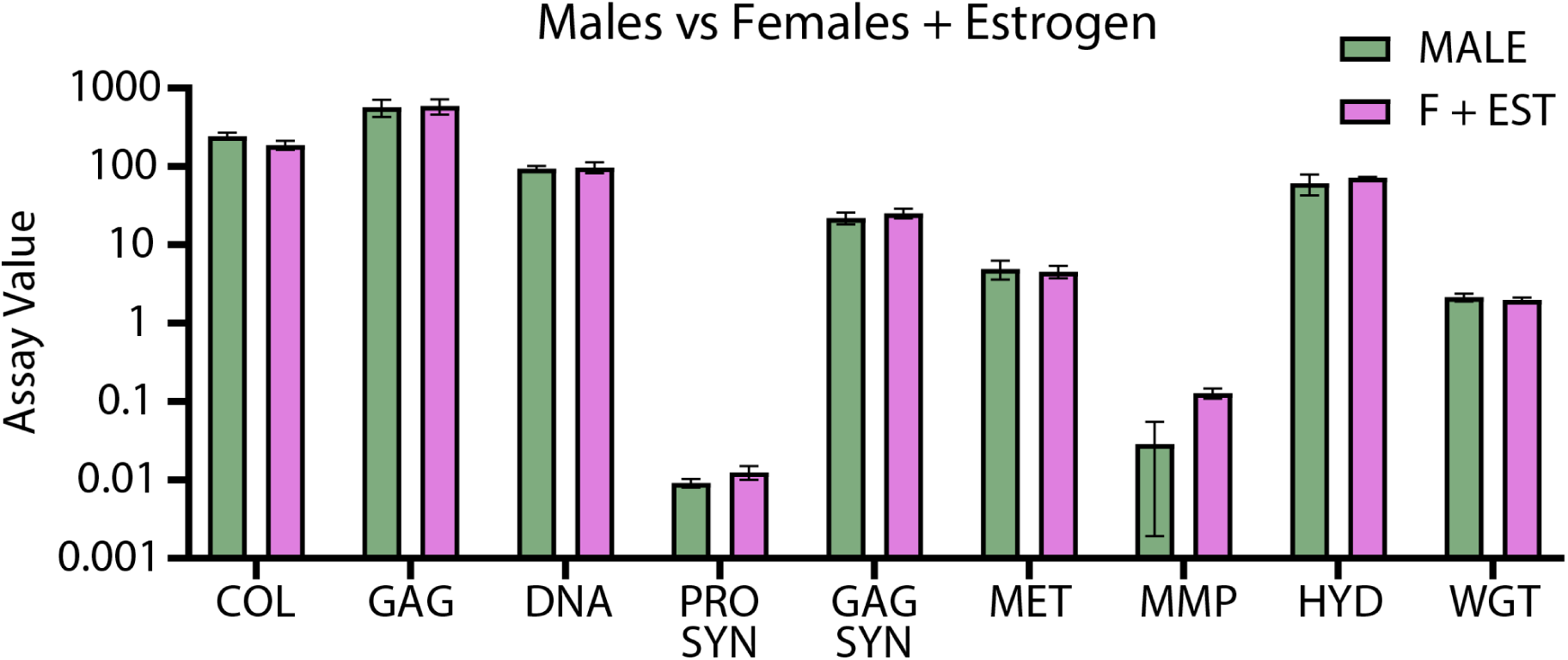
Comparison between male explants without hormone induction and female explants supplemented with estrogen after 7 days in culture. No significant differences are present between groups.

Progesterone seems to play a substantial role in female collagen turnover, increasing synthesis levels above that of males and increasing MMP activity. One interesting result is that a decrease in DNA content is preserved across timepoint and sex in response to progesterone, while explant metabolism remains constant. Viability and dose-dependency studies should be performed to further investigate this result. As with estrogen, progesterone has a greater influence on female ECM remodeling than in male tendons. The decreases in GAG and protein synthesis in males in response to progesterone may negatively impact tendon remodeling, but this is difficult to interpret without a measure of baseline synthesis parameters since these assays cannot be performed prior to culture. Lower synthesis rates are considered more physiologic, but in the context of injury repair, higher rates may be advantageous. Overall, progesterone seemed to facilitate the maintenance of baseline attributes. This could be extended to progesterone’s role in the maturation of tissues, as it had a large role in maintaining the incorporation of both collagen and GAGs, as seen in the content measurements. However, further mechanical and histological analysis is necessary to make this claim.

Interestingly, the dual hormone condition led to the most changes from both innate (day 0) and no hormone condition properties across both sexes. However, the impacts of this treatment were not consistently additive compared to individual hormone groups, suggesting the interplay between estrogen and progesterone is complex and assay-dependent. Degradative MMP activity and all measured gene expression were additive, while the explant health, biosynthesis, and content parameters were not. This implies that the individual hormones may have dominant impacts in specific cell processes. Interestingly, the dual hormone condition decreased GAG and protein turnover rates to levels more representative of a healthy physiologic state. Constant administration of single and dual hormones is not physiologically accurate, and these hormones are often present simultaneously at different concentrations. Therefore, it would be prudent to continue investigating the interplay between these hormones, as well as dose- and time-dependent effects of exogenous administration. This is also further motivation to explore the impacts that other important gonadotrophic hormones may have on tendon remodeling.

The addition of hormones led to *sex-dependent* changes in ECM turnover. Hormonal stimulation had a greater impact in female tissues than in males, leading to improved response to stress deprivation in females, but decreased responses in males. This is an important finding, as many studies in the literature use male models to assess the impact of female-associated sex hormones.^28,33,42,43^ Previous work exploring the response of male tissues to hormones in the context of acute injury suggest that estrogen is required for positive healing outcomes. In contrast, our work seems to indicate that hormones at supraphysiologic concentrations may actually impede the innate male response to injury. These data could have important implications for patients undergoing gender-affirming hormone therapies and supports more work needs to be done with respect to transgender health.

This study was not without limitations. First, hormonal treatments were supraphysiologic as they were chosen based on human serum levels during the menstrual cycle and levels used in murine cell culture.^29,42,49–53^ Therefore, they may not necessarily reflect the levels of hormones present in the mouse flexor tendon. Studies on hormone levels throughout the murine estrous cycle are limited, measured levels vary dramatically in both timing and concentration between studies, and there are no local tissue level studies at present. This is a significant knowledge gap that will need to be addressed as this body of work expands. In addition, residual hormones present in the FBS^63,64^ and phenol red^65–68^ in the culture medium have been shown to act like estrogen in some studies, and therefore may have interfered with our results. We are currently working to validate these studies further in phenol red-free and charcoal stripped medium.

Preliminary results suggest that hormones present in the medium may be interacting with other growth factors, bolstering matrix turnover. However, it is not yet clear that a filtering process like charcoal stripping would not remove other essential components, like IGF-1, which promotes healthy tendon metabolism. Chromosomal factors, which may contribute to sex differences through alterations of cell programming, could not be decoupled in this study design. Thus, we are working on including chromosomal invariant mice in future studies. Finally, stress deprivation responses only represent underuse injuries, not the more clinically relevant acute or chronic overuse injuries which may elicit different responses in the tendon. Other injury and dynamic loading models should be investigated, including microdamage and bulk mechanical injuries, extended underuse injuries, and dynamic loading scenarios mimicking homeostasis, exercise, and overuse.

Through the completion of this work, we aimed to determine the roles of sex and sex hormones on ECM remodeling for tendon maintenance and health. We revealed that males tend to rebuild matrix in response to stress deprivation while females better preserve baseline matrix properties. Hormones caused alterations to these innate responses, leading to sex-dependent changes in turnover activity. Overall, and in agreement with our hypothesis, estrogen seemed to have a dominant role in the upregulation of overall matrix turnover while progesterone was dominant in maintenance. However, there was also significant interaction between the two hormone treatments. Estrogen and progesterone, and the sex-dependent responses to each hormone, are thus critical to male and female tendon health and warrant further mechanistic studies regarding their effect on mechanosensing, healing, and adaptation.

## Acknowledgements

This study was supported by the National Institutes of Health (R35-GM151127) and an Agility Project Grant from the Wu Tsai Human Performance Alliance. The authors would like to also acknowledge Caitlin Colicchio for her contributions to discussions around this work.

## Author Contributions Statement

AS and BC contributed significantly to all aspects of the manuscript, including research design, the acquisition, analysis and interpretation of data, and manuscript preparation and editing. All authors have read and approved the final submitted manuscript.

